# RNA sequencing reveals interacting key determinants of osteoarthritis acting in subchondral bone and articular cartilage

**DOI:** 10.1101/2020.03.13.969386

**Authors:** Margo Tuerlings, Marcella van Hoolwerff, Evelyn Houtman, Eka (H.E.D.) Suchiman, Nico Lakenberg, Hailiang Mei, Enrike (H.M.J.) van der Linden, Rob (R.G.H.H.) Nelissen, Yolande (Y.F.M.) Ramos, Rodrigo Coutinho de Almeida, Ingrid Meulenbelt

**Author notes:** **Address for correspondence and reprint to** Ingrid Meulenbelt PhD, Department of Medical Statistics and Bioinformatics, Section of Molecular Epidemiology, Leiden University Medical Center, LUMC Post-zone S-05-P, PO Box 9600, 2300 RC Leiden, The Netherlands.

## Abstract

**Objective:** The aim of this study was to identify key determinants of the interactive osteoarthritis (OA) pathophysiological processes of subchondral bone and cartilage.

**Methods:** We performed RNA sequencing on macroscopically preserved and lesioned OA subchondral bone of patients that underwent joint replacement surgery due to OA (N=24 pairs; 6 hips, 18 knees, RAAK-study). Unsupervised hierarchical clustering and differential expression analyses were performed. Results were combined with previously identified, differentially expressed genes in cartilage (partly overlapping samples) as well as with recently identified OA risk genes.

**Results:** We identified 1569 genes significantly differentially expressed between lesioned and preserved subchondral bone, including *CNTNAP2* (FC=2.4, FDR=3.36×10^−5^) and *STMN2* (FC=9.6, FDR=3.36×10^−3^). Among the identified genes, 305 were also differentially expressed and with same direction of effects in cartilage, including *IL11* and *CHADL*, recently acknowledged OA susceptibility genes. Upon differential expression analysis stratifying for joint site, we identified 509 genes exclusively differentially expressed in subchondral bone of the knee, such as *KLF11* and *WNT4*. These exclusive knee genes were enriched for involvement in epigenetic processes, characterized by for instance *HIST1H3J* and *HIST1H3H*.

**Conclusion:** To our knowledge, we are the first to report on differential gene expression patterns of paired OA subchondral bone tissue using RNA sequencing. Among the most consistently differentially expressed genes with OA pathophysiology in both bone and cartilage were *IL11* and *CHADL.* As these genes were recently also identified as robust OA risk genes they classify as attractive druggable targets acting on two OA disease relevant tissues.

## INTRODUCTION

Osteoarthritis (OA) represents multiple subtypes of degenerative joint diseases, characterized by progressive and irreversible degeneration of the articular cartilage and structural changes in the subchondral bone. Globally, osteoarthritis is a highly prevalent and disabling disease which results in high social and economic burden to society [1]. Yet, there is no proven therapy to prevent or to slow down OA. Development of OA is dependent on multiple factors, with both environmental and genetic components [2, 3]. To discover genes and underlying disease pathways genetic studies, such as large genome wide association studies (GWAS), have been performed that identified compelling OA risk single nucleotide polymorphisms (SNPs) [4–6]. Functional follow-up studies involve exploring the expression patterns in disease relevant tissues, behavior with pathophysiology, and/or eQTL or cis-eQTL analysis. To date, major efforts have been made trying to characterize pathophysiological processes of OA in articular cartilage. However, only few studies focus on OA pathophysiological processes in the underlying bone [7, 8].

In the last decades, evidence has been accumulated that subchondral bone contributes to both onset and progression of OA [9–12]. In healthy bone, there is a balanced process between bone resorption and bone deposition as a consequence of dynamic adaptation to mechanical load. With OA, this balanced process is disturbed, which results in changes in the architecture of the subchondral trabecular bone, increased thickness of the subchondral bone plate, formation of new bony structures at the joint margins known as osteophytes, and development of subchondral bone cysts [2, 13, 14]. In addition, studies have shown an association between the bone mineral density and development of OA, which suggest that the subchondral bone is involved in the early stages of OA [13, 15]. This was also suggested by studies regarding subchondral bone marrow lesions, showing these to be very early markers of OA [8, 16].

In contrast to cartilage and despite its relevance, only a limited number of studies have been focusing on the characterization of OA disease processes on a gene expression level in the subchondral bone. Chou et al. [7] performed whole genome expression profiling of non-OA and OA subchondral bone using microarray analysis, which led to identification of genes involved in pathways such as lipid metabolism and mineral metabolism. Kuttapitiya et al. [8] used microarray analysis to identify genes involved in bone remodeling, pain sensitization, and matrix turnover being differentially expressed between OA bone marrow lesion tissue and controls. Nevertheless, these studies were performed using microarray and did solely include knee samples.

In this study, we explored RNA sequencing data of preserved and lesioned OA subchondral bone to identify genes that change with progression of OA. The samples used in this study were obtained from joints of patients that underwent total joint replacement surgery due to OA, as a part of the Research Arthritis and Articular Cartilage (RAAK) study. In total, we compared paired subchondral bone samples (preserved and lesioned) of 24 OA patients, of which also lesioned and preserved cartilage was collected. The results presented here, contribute to further understanding of the ongoing OA process in the subchondral bone and give insight into the pathophysiological processes in bone relative to cartilage.

## MATERIALS AND METHODS

### Sample description

The current study includes N=26 participants of the RAAK study, who underwent a joint replacement surgery as a consequence of OA. Macroscopically preserved and lesioned subchondral bone were collected from the joint of all patients and RNA sequencing was performed. Informed consent was obtained from all participants of the RAAK study and ethical approval for the RAAK study was given by the medical ethics committee of the Leiden University Medical Center (P08.239/P19.013).

### RNA sequencing

RNA was isolated from the subchondral bone using Qiagen RNeasy Mini Kit (Qiagen, GmbH, Hilden, Germany). Paired-end 2×100 bp RNA-sequencing (Illumina TruSeq RNA Library Prep Kit, Illumina HiSeq2000 and Illumina HiSeq4000) was performed. Strand specific RNA-seq libraries were generated which yielded a mean of 20 million reads per sample. Data from both Illumina platforms were integrated and analyzed with the same in-house pipeline. RNA-seq reads were aligned using GSNAP [17] against GRCh38 using default parameters. Read abundances per sample was estimated using HTSeq count v0.11.1 [18]. Only uniquely mapping reads were used for estimating expression. The quality of the raw reads for RNA-sequencing was checked using MultiQC v1.7. [19] The adaptors were clipped using Cutadapt v1.1 [20] applying default settings (min overlap 3, min length). To identify outliers, principal component analysis (PCA) was applied. After quality control only N=24 participants were included in further analysis.

### Cluster analysis

Prior to the cluster analysis, variance stabilizing transformation was performed on the data and 1000 genes were selected based on the highest coefficient of variation [21]. To identify the optimal number of clusters in the unsupervised hierarchical clustering the silhouette width score approach was used, with a higher average silhouette width score indicating a more optimal number of clusters [22].

### Differential expression analysis and pathway enrichment

Differential expression analysis was performed on paired lesioned and preserved subchondral bone samples, using the DESeq2 R package version 1.24.0 [23]. A general linear model assuming a negative binomial distribution was applied, followed by a paired Wald-test between lesioned and preserved OA samples, where the preserved samples were set as a reference. The Benjamini-Hochberg method was used to correct for multiple testing, as indicated by the false discorvery rate (FDR), with a significance cut-off value of 0.05. Gene enrichment was performed using the online functional annotation tool DAVID, selecting for the gene ontology terms Biological Processes (GOTERM_BP_DIRECT), Cellular Component (GOTERM_CC_DIRECT) and Molecular Function (GOTERM_MF_DIRECT) and for the Reactome and the KEGG pathways [24]. Moreover, the protein-protein interactions were analyzed using the online tool STRING version 11.0 [25]. An analysis summary scheme is shown in **Figure 1**.

**Figure 1.**
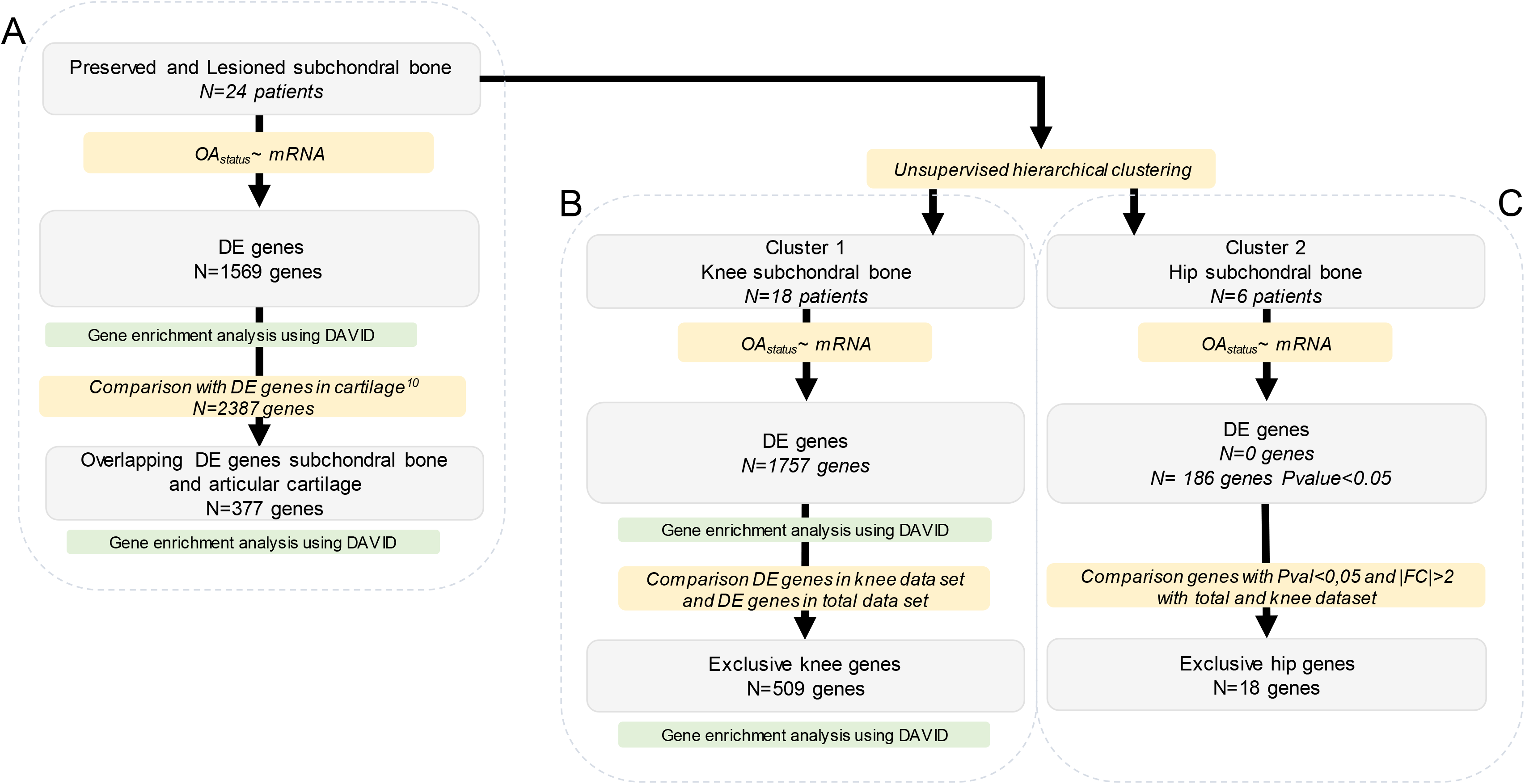
Overview of applied strategy. Number of genes represents the FDR significant differentially expressed (DE) genes, except for the hip genes.

### RT-qPCR validation

cDNA synthesis was done using Transcriptor First Strand cDNA Synthesis Kit (Roche, Basel, Switzerland), using 400 ng of RNA. RT-qPCR was performed to quantitatively determine gene expression of *FRZB, CNTNAP2, STMN2, CHRDL2, POSTN*, and *ASPN.* The relative gene expression was evaluated by the -ΔCT values, using *GAPDH* and *SDHA* as internal controls.

Generalized estimating equation (GEE) analysis was performed to calculate the statistical difference between the lesioned and preserved samples.

## RESULTS

### Sample characteristics

To characterize the pathophysiological process in subchondral bone with ongoing OA, we performed RNA sequencing on macroscopically preserved and lesioned OA subchondral bone samples of patients that underwent a joint replacement surgery due to OA (RAAK-study). The RNA-seq included N=24 paired samples (6 hips and 18 knees, **Supplementary Table 1**).

**Table 1.**
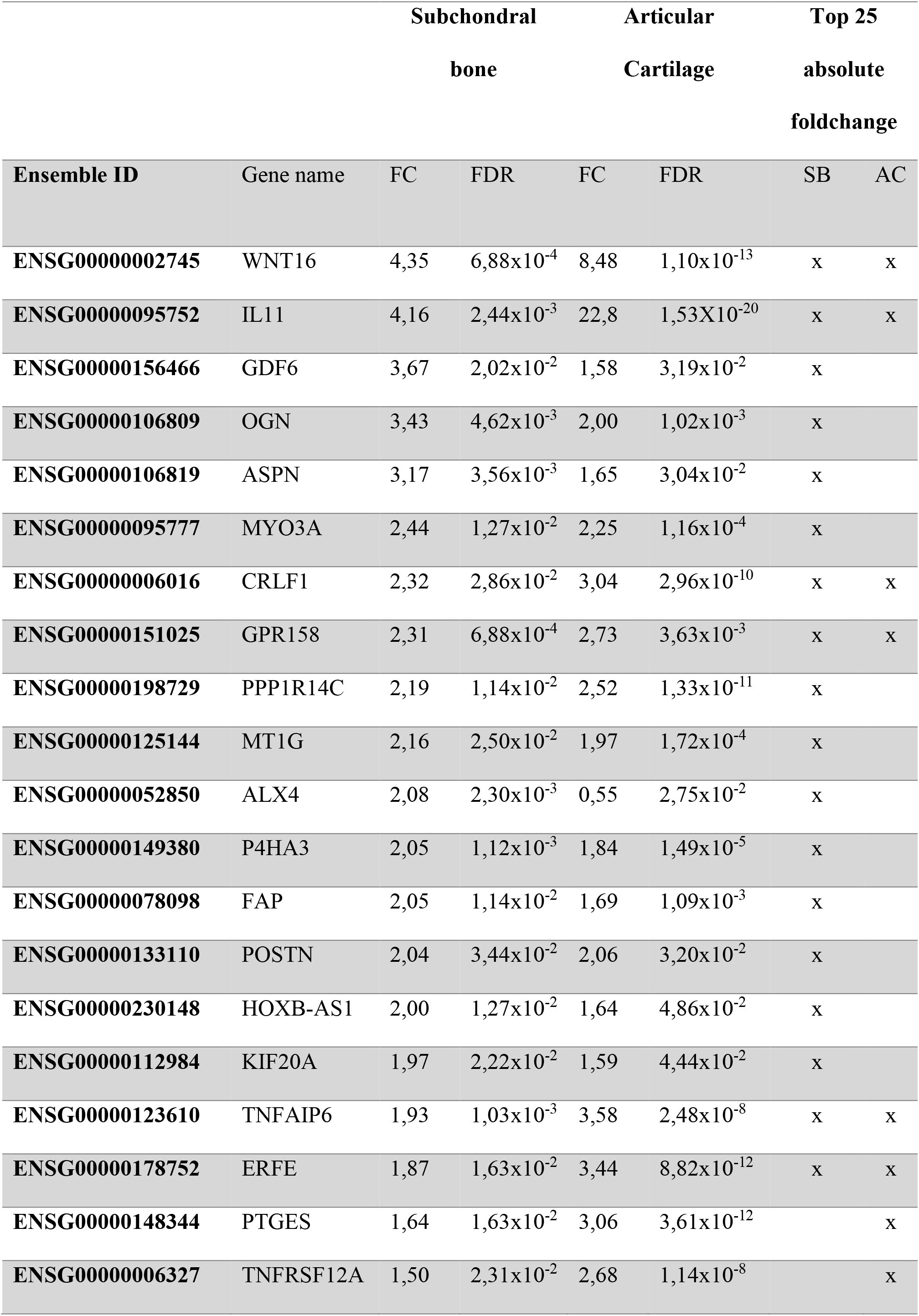
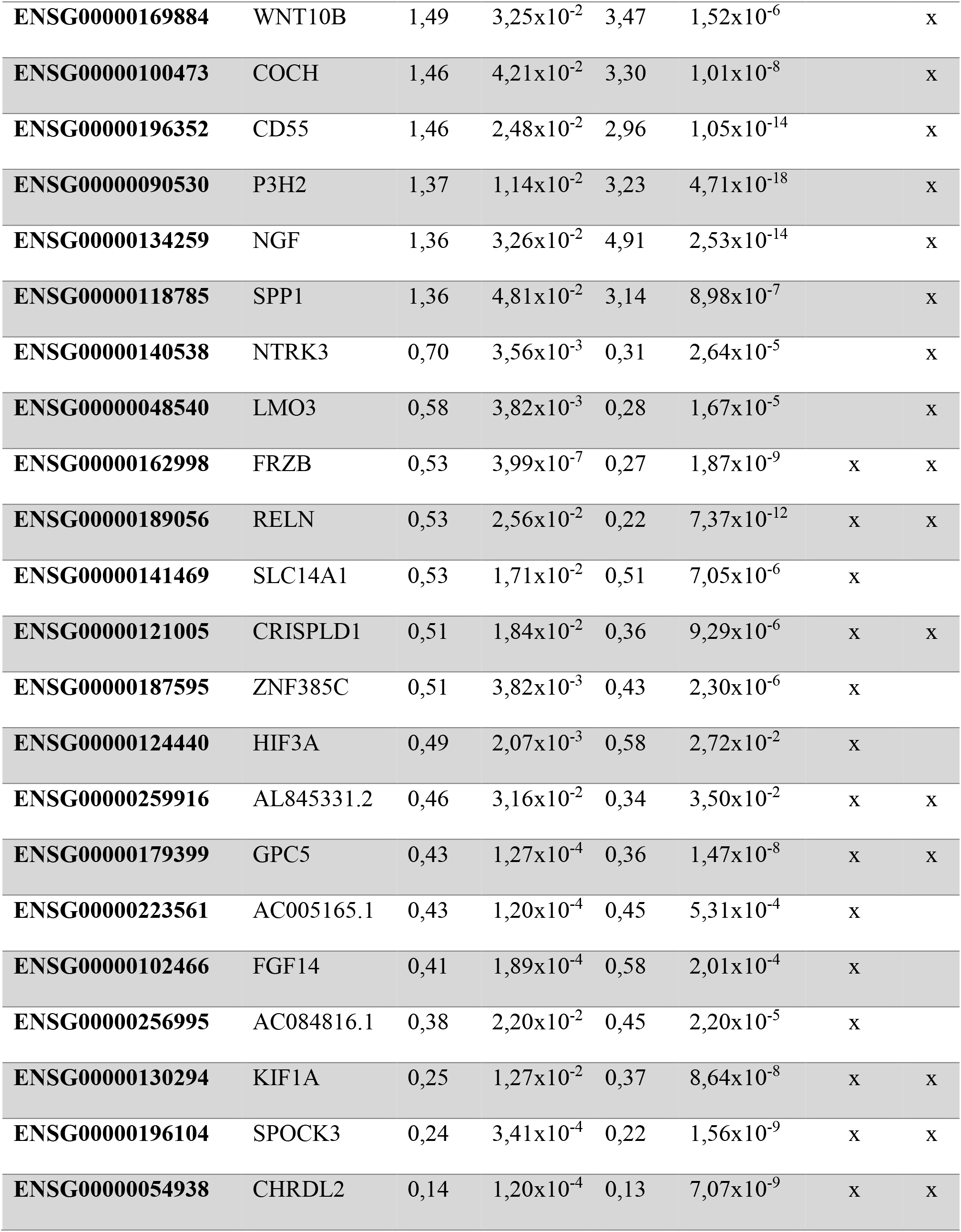
Genes that belonged to the top 25 genes based on the highest absolute foldchange in either bone or cartilage. Of these genes, 14 appear to be in the top 25 highest FC genes in both tissues.

Prior to the differential expression analysis, we tested possible contamination of cartilage in the subchondral bone samples. We used RNA-seq data of both tissues from the same joint to evaluate the expression levels of three cartilage specific genes and three bone specific genes (**Supplementary Table 2**) [26]. We observed significant lower expression levels of the cartilage markers and higher expression levels of the bone markers in the subchondral bone compared to the cartilage samples, indicating no to minimal contamination. Next, we explored whether the expression pattern in subchondral bone was associated to any baseline characteristics of patients (**Supplementary Table 1**), by performing unsupervised hierarchical clustering. To include the most informative genes to the cluster analysis, 1000 genes were selected based on the highest coefficient of variation in the total data set (preserved and lesioned, N=24 pairs). As shown in **Figure 2 (Supplementary Figure 1)**, we identified two clusters. These appeared to be based on joint site, indicating an inherent difference between hip and knee subchondral bone.

**Table 2.**
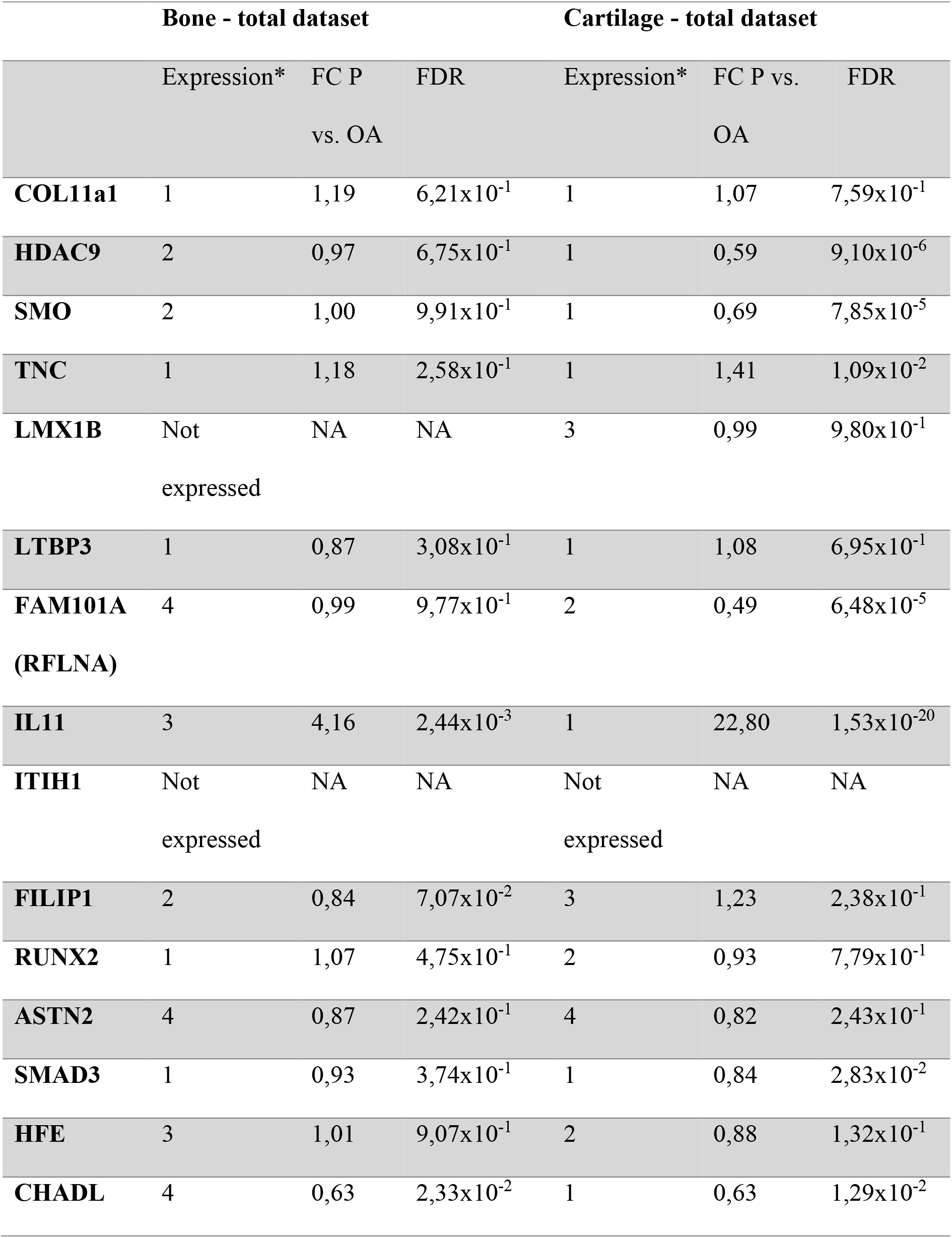
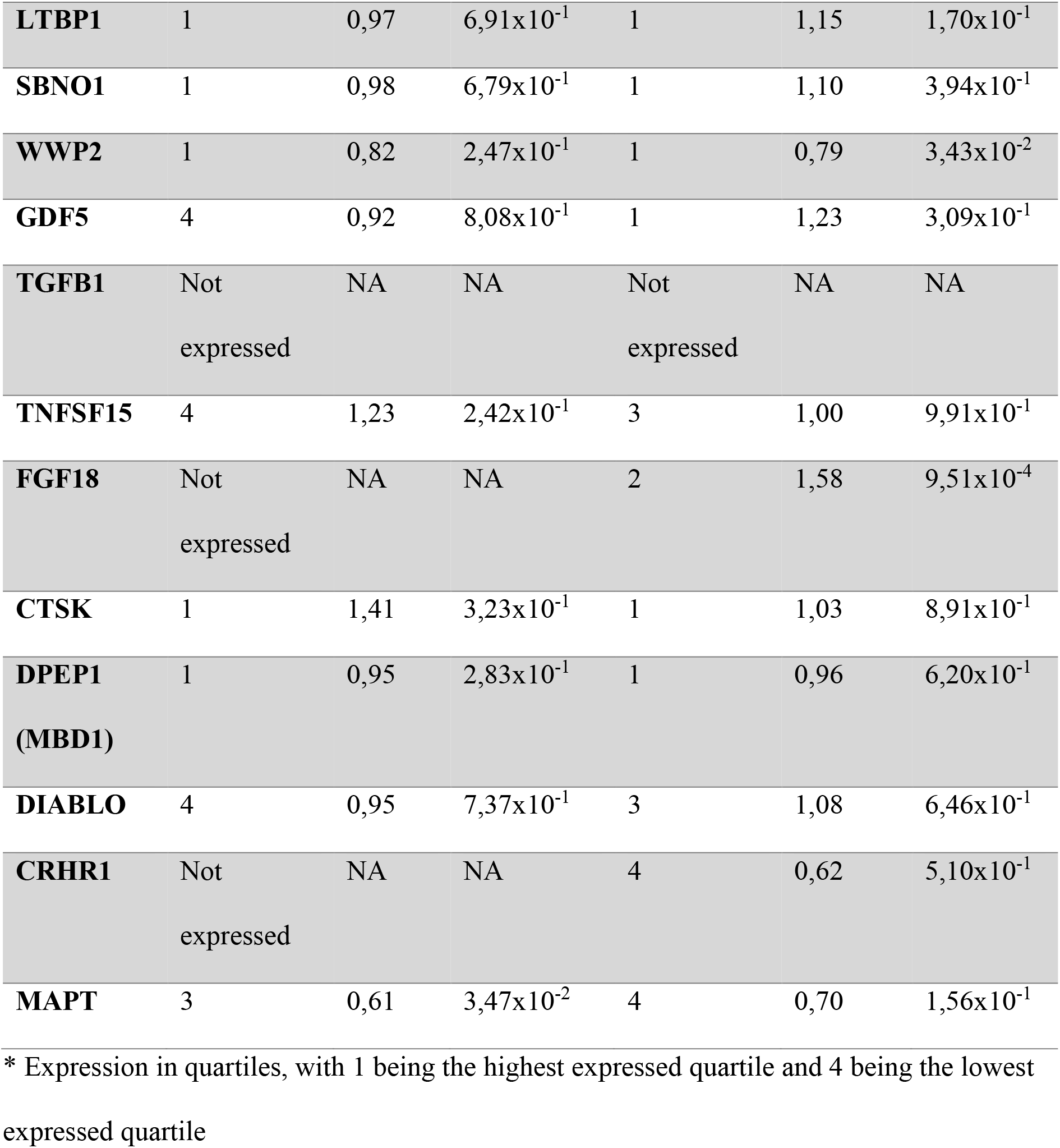
Expression levels and differential expression of new risk genes reported in two recent GWAS.

| Bone - total dataset |  |  |  | Cartilage - total dataset |  |  |
| --- | --- | --- | --- | --- | --- | --- |
|  | Expression* | FC P | FDR | Expression* | FC P vs. | FDR |
|  |  | vs. OA |  |  | OA |  |
| **COL11a1** | 1 | 1,19 | $6,21 \times 10^{-1}$ | 1 | 1,07 | $7,59 \times 10^{-1}$ |
| **HDAC9** | 2 | 0,97 | $6,75 \times 10^{-1}$ | 1 | 0,59 | $9,10 \times 10^{-6}$ |
| **SMO** | 2 | 1,00 | $9,91 \times 10^{-1}$ | 1 | 0,69 | $7,85 \times 10^{-5}$ |
| **TNC** | 1 | 1,18 | $2,58 \times 10^{-1}$ | 1 | 1,41 | $1,09 \times 10^{-2}$ |
| **LMX1B** | Not<br>expressed | NA | NA | 3 | 0,99 | $9,80 \times 10^{-1}$ |
| **LTBP3** | 1 | 0,87 | $3,08 \times 10^{-1}$ | 1 | 1,08 | $6,95 \times 10^{-1}$ |
| **FAM101A** | 4 | 0,99 | $9,77 \times 10^{-1}$ | 2 | 0,49 | $6,48 \times 10^{-5}$ |
| **(RFLNA)** |  |  |  |  |  |  |
| **IL11** | 3 | 4,16 | $2,44 \times 10^{-3}$ | 1 | 22,80 | $1,53 \times 10^{-20}$ |
| **ITIH1** | Not<br>expressed | NA | NA | Not<br>expressed | NA | NA |
| **FILIP1** | 2 | 0,84 | $7,07 \times 10^{-2}$ | 3 | 1,23 | $2,38 \times 10^{-1}$ |
| **RUNX2** | 1 | 1,07 | $4,75 \times 10^{-1}$ | 2 | 0,93 | $7,79 \times 10^{-1}$ |
| **ASTN2** | 4 | 0,87 | $2,42 \times 10^{-1}$ | 4 | 0,82 | $2,43 \times 10^{-1}$ |
| **SMAD3** | 1 | 0,93 | $3,74 \times 10^{-1}$ | 1 | 0,84 | $2,83 \times 10^{-2}$ |
| **HFE** | 3 | 1,01 | $9,07 \times 10^{-1}$ | 2 | 0,88 | $1,32 \times 10^{-1}$ |
| **CHADL** | 4 | 0,63 | $2,33 \times 10^{-2}$ | 1 | 0,63 | $1,29 \times 10^{-2}$ |
| **LTBP1** | 1 | 0,97 | $6,91 \times 10^{-1}$ | 1 | 1,15 | $1,70 \times 10^{-1}$ |
| **SBNO1** | 1 | 0,98 | $6,79 \times 10^{-1}$ | 1 | 1,10 | $3,94 \times 10^{-1}$ |
| **WWP2** | 1 | 0,82 | $2,47 \times 10^{-1}$ | 1 | 0,79 | $3,43 \times 10^{-2}$ |
| **GDF5** | 4 | 0,92 | $8,08 \times 10^{-1}$ | 1 | 1,23 | $3,09 \times 10^{-1}$ |
| **TGFB1** | Not<br>expressed | NA | NA | Not<br>expressed | NA | NA |
| **TNFSF15** | 4 | 1,23 | $2,42 \times 10^{-1}$ | 3 | 1,00 | $9,91 \times 10^{-1}$ |
| **FGF18** | Not<br>expressed | NA | NA | 2 | 1,58 | $9,51 \times 10^{-4}$ |
| **CTSK** | 1 | 1,41 | $3,23 \times 10^{-1}$ | 1 | 1,03 | $8,91 \times 10^{-1}$ |
| **DPEP1**<br>**(MBD1)** | 1 | 0,95 | $2,83 \times 10^{-1}$ | 1 | 0,96 | $6,20 \times 10^{-1}$ |
| **DIABLO** | 4 | 0,95 | $7,37 \times 10^{-1}$ | 3 | 1,08 | $6,46 \times 10^{-1}$ |
| **CRHR1** | Not<br>expressed | NA | NA | 4 | 0,62 | $5,10 \times 10^{-1}$ |
| **MAPT** | 3 | 0,61 | $3,47 \times 10^{-2}$ | 4 | 0,70 | $1,56 \times 10^{-1}$ |

\* Expression in quartiles, with 1 being the highest expressed quartile and 4 being the lowest expressed quartile

**Figure 2.**
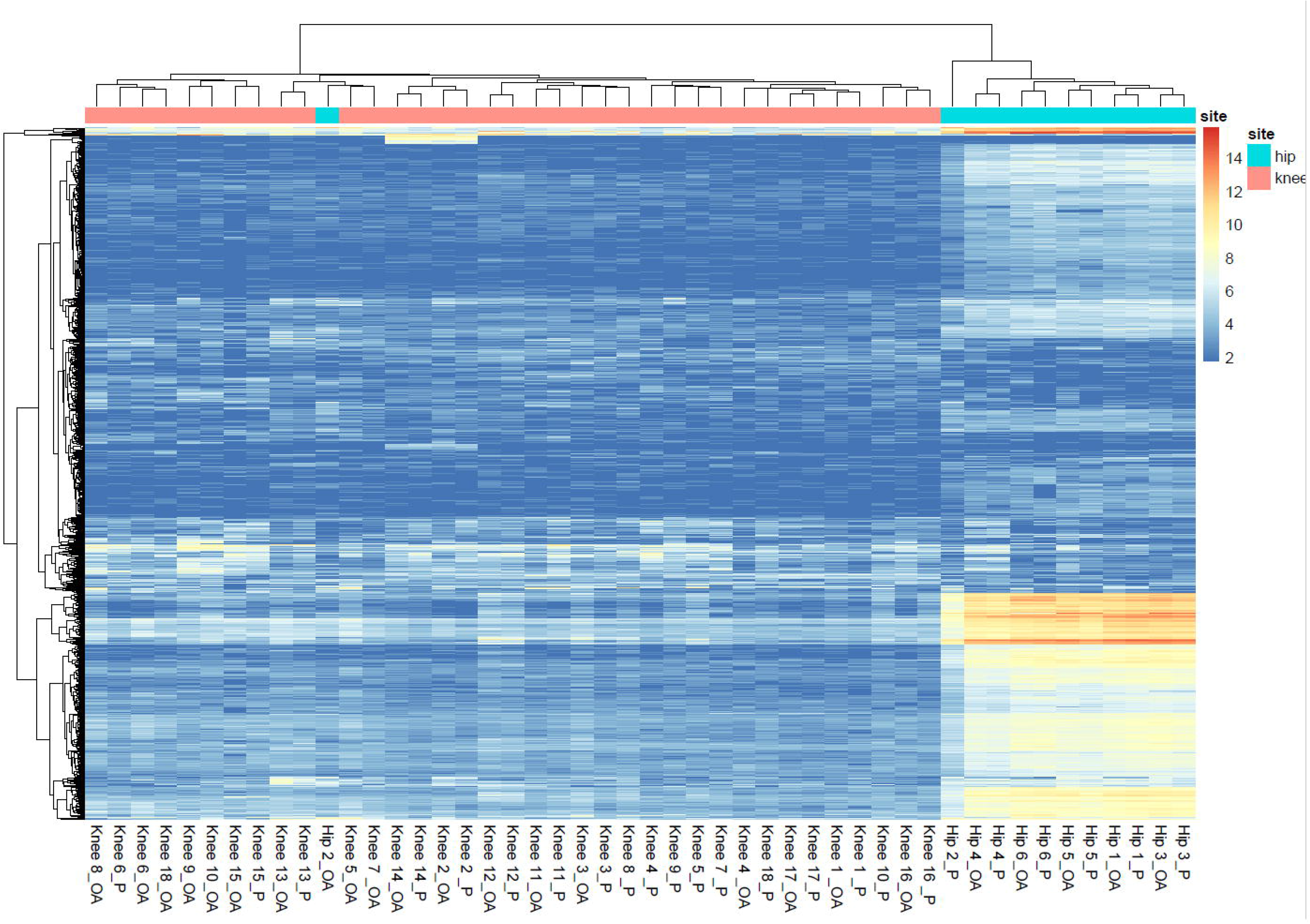
Cluster analysis based on the 1000 genes selected for their highest COV. Two clusters were identified based on knee samples (left) and hip samples (right).

### Differential expression analysis and pathway enrichment

We first determined the consistent differentially expressed (DE) genes between preserved and lesioned OA subchondral bone in the overall data set to explore the most consistent OA pathways (**Figure 1A**). Upon differential expression analysis (N=24 pairs), we identified 1569 genes being genome wide significantly DE between lesioned and preserved OA subchondral bone tissue. Of these DE genes, 750 were upregulated and 819 were downregulated (**Figure 3, Supplementary Table 3**). The most significantly downregulated gene was *FRZB* (FC=0.53, FDR=3.99×10^−7^), encoding the frizzled receptor protein, which is a well-known OA gene showing consistent lower expression in lesioned relative to preserved OA articular cartilage [27, 28]. The most significantly upregulated gene was *CNTNAP2* (FC=2.42, FDR=3.36×10^−5^), encoding the contactin-associated protein-like 2 protein (CASPR2). Among the 1569 DE genes, 53 genes had an absolute foldchange of 2 or higher (35 upregulated and 18 downregulated). The highest upregulated gene was *STMN2* (FC=9.56, FDR=2.36×10^−3^), encoding stathmin 2, while the most downregulated gene was *CHRDL2* (FC=0.14, FDR=1.20×10^−4^), encoding chordin-like protein 2.

**Figure 3.**
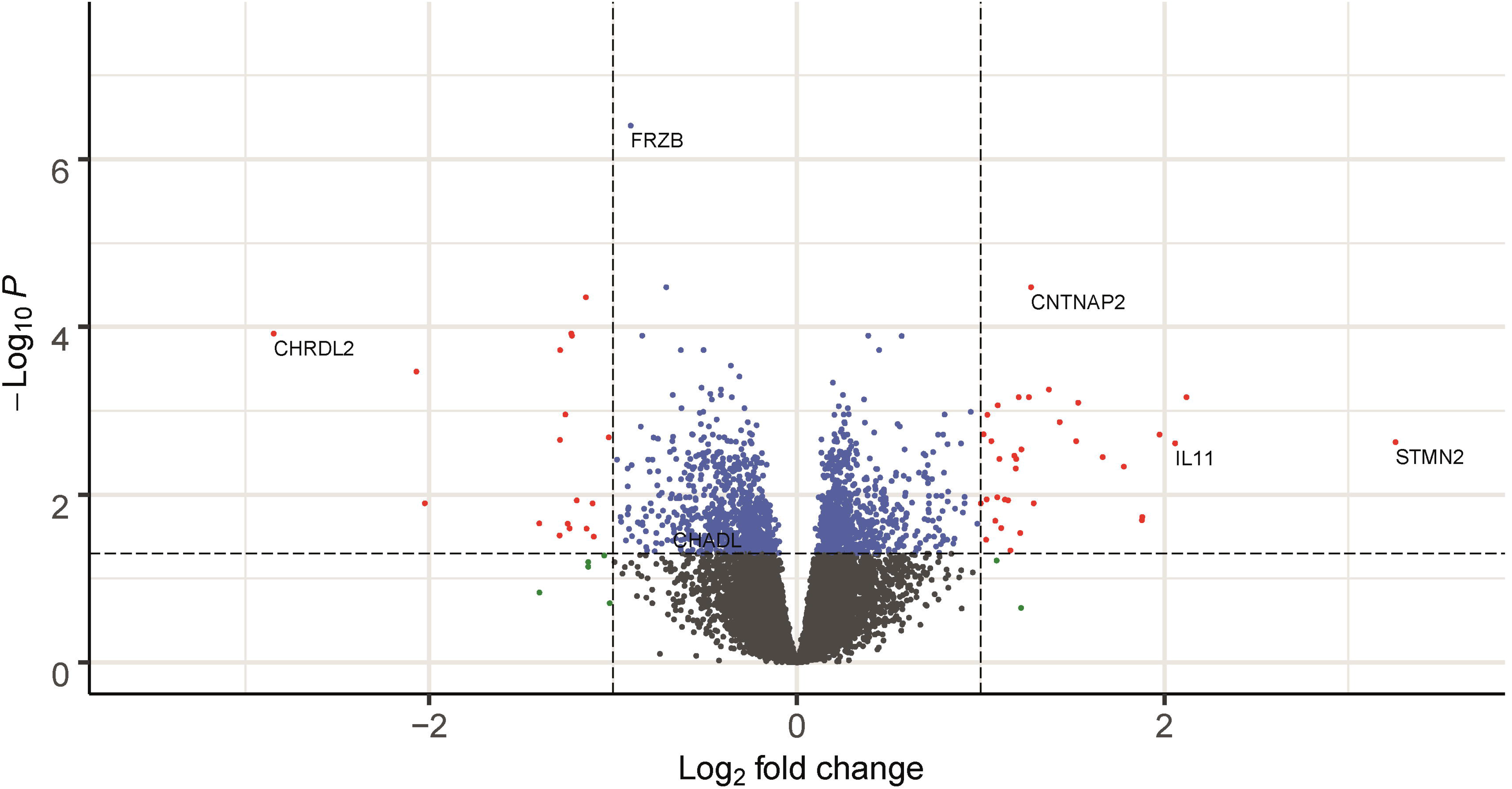
Volcano plot of differential expressed genes in the subchondral bone. The dots in de figure represent genes expressed in bone. Blue dots represent genes that are significantly differentially expressed, red dots represent genes that are significantly differentially expressed and have an absolute foldchange of 2 or higher, and green dots represent the genes with an absolute fold change of two or higher that are not significantly differentially expressed.

Next, we explored whether the significant DE genes (N=1569 genes) were enriched to particular pathways or processes using DAVID. Results indicated significant enriched GO-terms regarding processes involved in translational and post-translational processes, such as SRP-dependent co-translational protein targeting to membrane (GO:0006614, 33 genes, FDR=4.27×10^−7^) and translational initiation (GO:0006413, 36 genes, FDR=1.95×10^−4^). These processes were both mainly characterized by ribosomal proteins such as *RPS24*, *RPS4X* and *RPS18* (**Supplementary Table 4)**. Gene enrichment analysis of the genes selected for the highest FC (FC>|2|, N=53 genes), resulted in significant enrichment of processes regarding the extracellular matrix (GO:0005615, 16 genes, FDR=1.19×10^−5^), characterized by the upregulation of *WNT16* (FC=4.35, FDR= 6.88×10^−4^) *, CRLF1* (FC=2.32, FDR=2.86×10^−2^) and *OGN* (FC=3.43, FDR= 4.62×10^−3^), and the proteinaceous extracellular matrix (GO:0005578, 7 genes, FDR=4.50×10^−2^), characterized by upregulation of *POSTN* (FC=2.04, FDR=3.44×10^−2^), *ASPN* (FC=3.17, FDR=3.56×10^−3^) and *CTHRC1* (FC=2.15, FDR=3.75×10^−3^) (**Supplementary Table 5**). Furthermore, to explore interactions between proteins encoded by DE genes showing a FC>|2| (N=53genes) we used the STRING online tool. We identified a significant enrichment for protein-protein interactions (PPI) among 22 out of 44 proteins (P=3.20×10^−9^, **Figure 4**).

**Figure 4.**
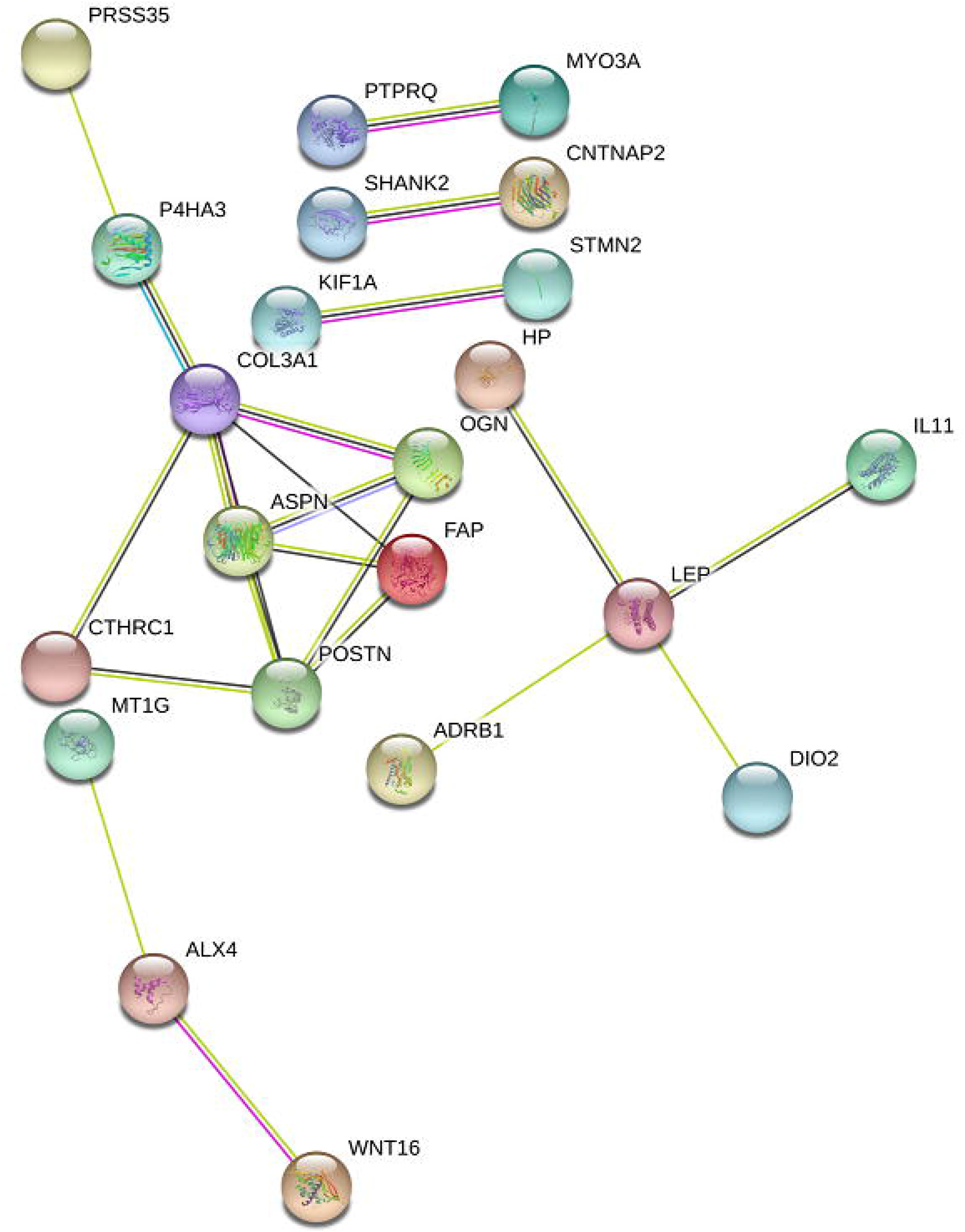
Protein-protein interaction network of proteins encoded by genes that show an absolute foldchange of 2 or higher (N=53 genes) created by STRING.

### Comparison subchondral bone and articular cartilage

To investigate interacting OA pathophysiological processes in subchondral bone and articular cartilage, we compared the identified DE genes in bone with our previously reported results on DE genes in articular cartilage [27] (**Figure 1A**, N=24 pairs for bone, N=35 pairs for cartilage, N=14 overlapping between bone and cartilage). As a result, we found an overlap of 337 genes being DE in both subchondral bone and articular cartilage (**Supplementary Figure 2**). Of these 337 overlapping genes, the majority (305 genes) showed similar direction of effect (**Supplementary Table 6**), while 32 genes showed an opposite direction of effect between both tissues (**Supplementary Table 7**). *ALX4,* encoding ALX homeobox 4, was a notable gene among the genes showing opposite direction of effects. ALX is known to be involved in osteogenesis and it is one of the highest upregulated genes in bone (**Table 1**). Among the 305 genes showing similar direction of effects, 14 genes belonged to the top 25 of genes with the highest foldchange in both tissues, such as *WNT16*, *IL11*, *CRLF1* and *FRZB* (**Table 1**). To explore common underlying pathways in the subchondral bone and the articular cartilage, we performed gene enrichment analysis with the 305 genes showing similar direction of effects. As a result, we found significant enrichment for the GO-terms extracellular region (GO:0005576, 36 genes, FDR= 4.56×10^−3^), characterized by the expression of for example *COL6A3*, *FGF14* and *GDF6*, proteinaceous extracellular matrix (GO:0005578, 17 genes, FDR= 7.98×10^−3^), characterized by the expression of for example *CHADL*, *ADAMTS17* and *SPOCK3,* and extracellular space (GO:0005615, 37 genes, FDR= 4.42×10^−3^), characterized by the expression of for example *CD63*, *SPP1* and *RELN* (**Supplementary Table 8**).

### Differential expression analysis stratified for joint site

Since hip and knee showed different gene expression profiles according to the cluster analysis (**Figure 2**), we repeated the differential expression analysis stratified for joint site to explore whether we could identify exclusive OA pathways that occur in subchondral bone of knees or hips only. Upon performing differential expression analysis on knee samples (N=18 pairs) as indicated by **Figure 1B**, we found 1757 genes that were significantly DE, of which 901 genes were upregulated and 855 genes were downregulated in lesioned compared to preserved OA subchondral bone (**Supplementary Table 9**). Moreover, 509 DE exclusive knee genes were identified (**Supplementary Table 10**); i.e. DE genes not present in the identified DE genes using the total dataset (**Supplementary Table 3**) nor using the hip dataset (**Supplementary Table 12**). Enrichment analysis of these exclusive knee genes showed significant enrichment for processes involved in epigenetic regulation, such as nucleosome (GO:0000786, 20 genes, 1.81×10^−9^), DNA methylation (R-HSA-5334118, 15 genes, 2.48×10^−6^) and regulation of gene silencing (GO:0060968, 6 genes, 1.90×10^−2^), all characterized by members of H3 histone family, such as *HIST1H3J* and *HIST1H3H* (**Supplementary Table 11).**

Differential expression analysis only with the hip samples (N=6 pairs) did not show any FDR significantly DE genes when comparing preserved and lesioned subchondral bone (**Figure 1C**). Nonetheless, among the genes with a P-value smaller than 0.05 and an absolute foldchange of 2 or higher (**Supplementary Table 12),**18 genes appeared to be exclusive for hip; i.e. genes not present in the identified DE genes using the total dataset (**Supplementary Table 3**) nor the knee dataset (**Supplementary Table 9**). Among these exclusive hip genes, we found *CALCR*, *LGR5* and *COL2A1* (**Supplementary Table 13**).

### Validation of differentially expressed genes

To validate and replicate the findings of the differential expression analysis on the RNA-seq data, we included a set of N=20 samples to perform both technical (N=10 samples) and biological (N=10 samples) replication by RT-qPCR. Validation and replication of six genes: *FRZB*, *CNTNAP2*, *STMN2*, *CHRDL2*, *POSTN*, and *ASPN* showed a significant difference between preserved and lesioned subchondral bone, with similar direction of effects. Replication also showed significant differences, with the same direction of effects as reported here for the RNA-seq data (**Supplementary Table 14**).

### Differential expression of previously identified risk genes

In recent genome-wide association studies (GWAS) of hip and knee OA [5, 6], 27 new loci conferring risk to OA were identified (**Table 2**). To see whether those OA susceptibility genes are sensitive to OA pathophysiology in either the articular cartilage, the subchondral bone or both, we explored the expression levels and the differential expression between lesioned and preserved tissue of those genes in our data sets. As shown in **Table 2**, we found two risk genes, *IL11* and *CHADL,* being differentially expressed in both the subchondral bone and the articular cartilage. In addition, *IL11* showed a significant differential expression in knee subchondral bone (FC=4.07, FDR=7.00×10^−3^) and showed a high foldchange (FC=4.77, Pval= 4,43×10^−02^) in hip subchondral bone. This indicates that, based on our data set, *IL11* has an effect in both tissues and in both joint sites, albeit not FDR significant in hip subchondral bone.

## DISCUSSION

Differential expression analysis of the gene expression levels between preserved and lesioned OA subchondral bone (N=24 paired samples), showed 1569 genes being significantly DE, including *CNTNAP2* and *STMN2*. Upon comparing these 1569 DE genes with the 2387 genes previously shown to be DE with OA pathophysiology in cartilage, we found an overlap of 305 genes that had the same direction of effect. These 305 overlapping genes were enriched for processes related to the extracellular matrix, characterized by the expression of amongst others *COL6A3, GDF6* and *SPP1*. Moreover, among the 305 overlapping genes were *IL11* and *CHADL* (**Supplementary Table 6**), which were previously identified as being genetic OA risk genes (**Table 2**). By applying hierarchical clustering on the overall RNA-seq dataset of the subchondral bone, we identified two clusters based on joint site (knees and hips). When stratifying the analysis for joint site, we identified 1759 genes DE between preserved and lesioned knee OA bone, of which 509 genes were exclusive knee genes, including genes such as *WNT4* and *KLF11*. These exclusive DE knee OA genes were, amongst others, enriched for regulation of gene silencing by epigenetic processes, such as DNA methylation and histone modification, characterized by genes such as *HIST1H3J* and *HIST1H3H*.

Among the 1569 genes being FDR significantly DE between lesioned and preserved OA subchondral bone in the complete dataset, we identified *CNTNAP2* (FC=2.42, FDR=3.36×10^−5^) and *STMN2* (FC=9.56, FDR=2.36×10^−3^) as the most significantly upregulated gene and the gene with the highest FC, respectively. *CNTNAP2*, encoding CASPR2, is known for its effect on cell-cell interactions in the nervous system, synapse development, neural migration and neural connectivity [29, 30]. Neither *CNTNAP2* or its encoded protein were previously identified as being related to OA. *STMN2* also plays a role in the control of neuronal differentiation. Moreover, *STMN2* is expressed during osteogenesis and it was previously shown to be highly upregulated in OA bone marrow lesions as compared to control bone samples [8, 31]. In addition, we found other neural markers to be upregulated in lesioned compared to preserved OA subchondral bone, such as *NGF* and *THBS3* (**Supplementary Table 3**). The upregulation of those genes indicate that the formation of new neuronal structures in bone is increased with ongoing OA. These data may confirm that OA related pain originates from bone [8].

Based on genes with the highest coefficient of variance, we identified two clusters representing knee and hip subchondral bone, respectively. These clusters indicate distinct inherent transcriptome landscapes between the two joint sites, which was not previously seen with similar analyses of the cartilage (unpublished data). Upon differential expression analysis stratifying for joint site, we discovered 509 unique knee genes compared to the complete dataset, which were significantly enriched for epigenetic processes such as DNA methylation reflected by the expression of among others *HIST1H3J* and *HIST1H3H*. The significant enrichment of these epigenetic processes amongst the unique knee genes indicates a change in epigenetics with ongoing knee OA, which is not seen with ongoing hip OA. This was also previously seen in the articular cartilage, where hip and knee methylation profiles cluster apart irrespective of the OA status. However, this was characterized by the expression of different genes, such as the HOX genes [32, 33]. We did not find FDR significant genes when selecting the hip samples, which is likely due to the small sample size (N=6 pairs). However, we found 18 exclusive genes DE in hip based on the nominal P-value and an absolute foldchange of 2 or higher, including genes such as *CALCR*, *LGR5* and *COL2A1*. However, replication is needed to confirm these exclusive hip genes.

Given the accumulating awareness of crosstalk between the articular cartilage and the subchondral bone during OA [10, 34], we here report on the comparison between RNA-seq data of the subchondral bone and the articular cartilage (N=24 pairs, N=35 pairs, respectively, with an overlap of N=14 patients). As reflected by the relatively small overlap in DE genes (9.28%, 305 genes) between subchondral bone and cartilage, there is a difference in OA pathophysiology between the two tissues.

To find genes that are most likely causal to OA, we explored 27 previously published genes in which single nucleotide polymorphisms (SNPs) were identified as being genome-wide significantly associated with OA (**Table 2**), suggesting that those genes have a more causal relationship to OA and making them attractive potential drug targets [5, 6]. To see whether the previously identified OA risk genes are involved in the OA pathophysiological process in both tissues, we compared the expression levels and the differential expression between the preserved and lesioned samples (**Table 2**). As a result, we found the OA risk genes *IL11* and *CHADL* being differentially expressed in both tissues and showing the same direction of effect and thus making them attractive potential drug targets with effects in both tissues. *CHADL*, encoding Chondroadherin-Like Protein, is involved in collagen binding and is a negative modulator of chondrocyte differentiation. The OA susceptibility allele rs117018441-T, located in an intron of *CHADL*, marks higher expression of *CHADL* compared to rs117018441-G in skeletal muscle and adipose tissue (GTEx) [5, 35]. This may indicate that increased expression of *CHADL* has a negative regulatory role both in bone and cartilage and that a therapeutic strategy could be the inhibition of this gene. However, when stratifying for joint site, we found *CHADL* being DE in particularly the knee subchondral bone, suggesting *CHADL* is a potential drug target for knee OA exclusively. *IL11*, encoding Interleukin 11, is known for its role in bone remodelling and lack of IL11 function is associated with impaired bone formation [36]. Notably, IL11 is recently proposed as potential therapeutic drug target for OA in cartilage [6], since the OA risk allele rs4252548-T, a missense variant p.Arg112His, acts via reduced function of the IL11 protein. As such, increasing IL11 protein levels was proposed as a therapeutic strategy for treatment of OA. We here again show that *IL11* is highly upregulated between preserved and lesioned OA tissue in both subchondral bone and articular cartilage (FC=4.16 and FC=22.8, respectively). Together these data indicate that reduced function of IL11 is predisposing to OA onset and that the upregulation of *IL11* with OA pathophysiology could be considered an attempt of the chondrocytes to enhance ECM integrity. Nonetheless, the consistent and considerable upregulation of *IL11* in both the subchondral bone and the articular cartilage may not necessarily a lack of potency to produce IL11, unless translation of the protein is hampered. This needs to be further functionally investigated preferably in an *in vitro* model of OA. Altogether, *CHADL* and *IL11* could both be a highly suitable drug target with effects in both bone and cartilage. However, further functional research is necessary to confirm the effects of *CHADL* and *IL11* on bone and cartilage metabolism.

To our knowledge, we are the first to report on the large scale differential expression pattern of OA subchondral bone using RNA sequencing for both hip and knee samples. We identified distinct expression patterns between hips and knees. Moreover, we identified multiple genes that were previously identified in OA articular cartilage and we identified genes that are subchondral bone specific. Together these data will contribute to a better understanding of pathophysiological process underlying development of OA.

## Supporting information

Supplementary Figure 1 - Silhouette width score showing an optimal number of two clusters.

Supplementary Figure 2 - Venn diagram of differentially expressed genes in the articular cartilage (N=2387) and in the subchondral bone (N=1569).

## ACKNOWLEDGEMENTS

We thank all the participants of the RAAK study. The LUMC has and is supporting the RAAK study. We also thank Demien Broekhuis, Robert vd Wal, Peter van Schie, Shaho Hasan, Maartje Meijer, Daisy Latijnhouwers and Geert Spierenburg for collecting the RAAK material. The study was funded by the Dutch Scientific Research council NWO /ZonMW VICI scheme (nr 91816631/528), Dutch Arthritis Society (DAA_10_1-402), and European Commission Seventh Framework programme (TreatOA, 200800). Moreover, we thank the Medical Delta. Finally, we thank the Sequence Analysis Support Core (SASC) of the Leiden University Medical Center for their support.

## FIGURE LEGENDS

**Supplementary Figure 1 – Silhouette width score showing an optimal number of two clusters.**

**Supplementary Figure 2 – Venn diagram of differentially expressed genes in the articular cartilage (N=2387) and in the subchondral bone (N=1569).** 337 genes were overlapping between cartilage and bone, of which 305 genes show similar direction of effects between cartilage and bone.

