## Supplementary figures and images for "RNA sequencing reveals interacting key determinants of osteoarthritis acting in subchondral bone and articular cartilage"

### Supplementary Figure 1 - Silhouette width score showing an optimal number of two clusters.

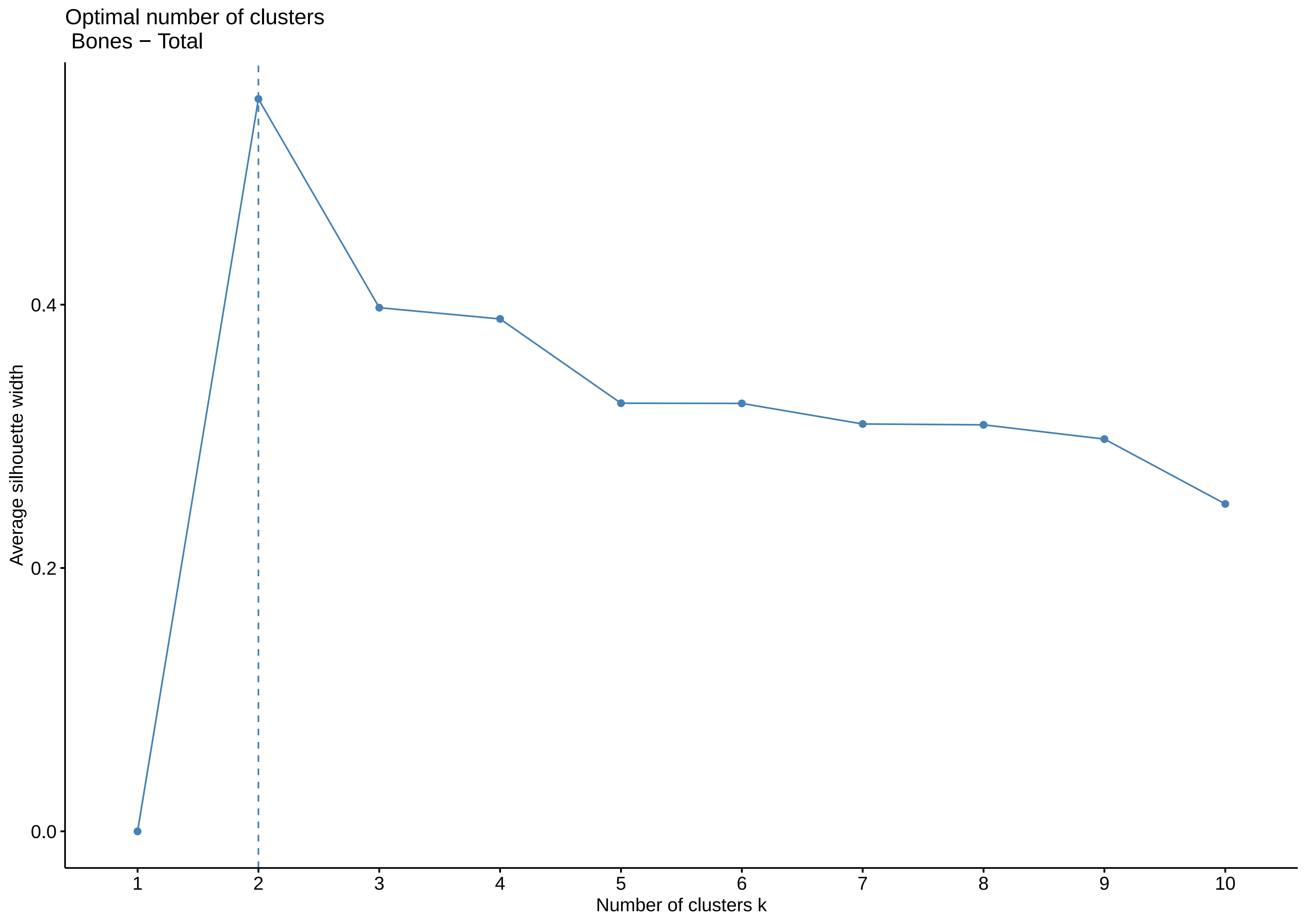

### Supplementary Figure 2 - Venn diagram of differentially expressed genes in the articular cartilage (N=2387) and in the subchondral bone (N=1569).

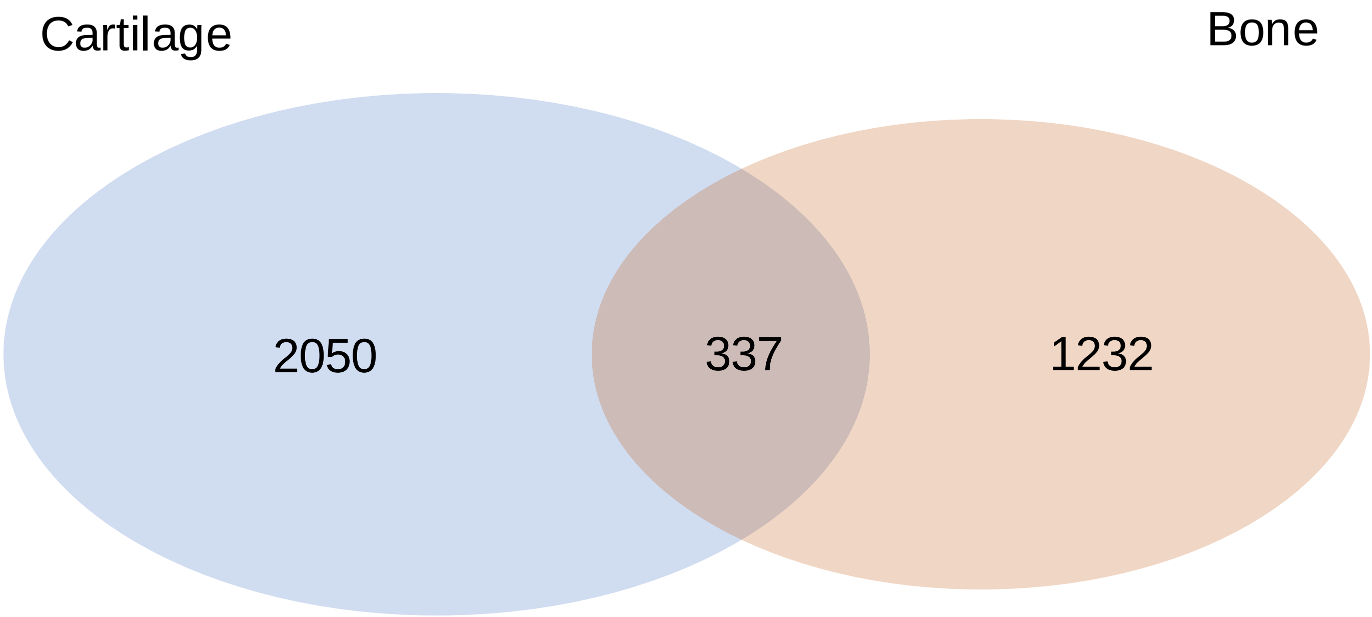
